# Analysis of Pan-Omics Data in Human Interactome Network (APODHIN)

**DOI:** 10.1101/2020.04.18.048207

**Authors:** Nupur Biswas, Krishna Kumar, Sarpita Bose, Raisa Bera, Saikat Chakrabarti

## Abstract

Analysis of Pan-Omics Data in Human Interactome Network (APODHIN) is a platform for integrative analysis of transcriptomics, proteomics, genomics, and metabolomics data for identification of key molecular players and their interconnections exemplified in cancer scenario. APODHIN works on a meta-interactome networks consisting of human protein-protein interactions, miRNA-target gene regulatory interactions, and transcription factor-target gene regulatory relationships, respectively. In its first module, APODHIN maps proteins/genes/miRNAs from different omics data in its meta-interactome network and extracts the network of biomolecules that are differentially altered in the given scenario. Using this context specific, filtered interaction network, APODHIN identifies topologically important nodes (TINs) implementing graph theory based network topology analysis and further justifies their role via pathway and disease marker mapping. These TINs could be used as prospective diagnostic and/or prognostic biomarkers and/or potential therapeutic targets. In its second module, APODHIN attempts to identify cross pathway regulatory and protein-protein interaction (PPI) links connecting signaling proteins, transcription factors, and miRNAs to metabolic enzymes via utilization of single-omics and/or pan-omics data and implementation of mathematical modeling. Interconnections between regulatory components such as signaling proteins/TFs/miRNAs and metabolic pathways need to be elucidated more elaborately in order to understand the role of oncogene and tumor suppressors in regulation of metabolic reprogramming during cancer.

APODHIN platform contains a web server component where users can upload single/multi omics data to identify TINs and cross-pathway links. Tabular, graphical and 3D network representations of the identified TINs and cross-pathway links are provided for better appreciation. Additionally, this platform also provides a database part where cancer specific, single and/or multi omics dataset centric meta-interactome networks, TINs, and cross-pathway links are provided for cervical, ovarian, and breast cancers, respectively. APODHIN platform is freely available at http://www.hpppi.iicb.res.in/APODHIN/home.html.

## Introduction

Technological advances have made different types of omics data accessible in large scale. Different types of omics data include RNA transcriptomics, miRNA transcriptomics, proteomics, phosphoproteomics, genomics, epigenomics, metabolomics, lipidomics, and pharmacogenomics which are also considered as ‘big’ data in the context of biological data analysis. Collective analysis of these multi-dimensional data is referred to as ‘pan-omics’ [1]. Integration of these pan-omics data is highly needed to interpret the underlying biological processes in the genesis and progression of systemic/genetic diseases. Because of the heterogeneous nature of the diseases, even if patients having similar symptoms are treated similarly, the disease prognosis differs a lot. It shows the inadequacy of symptom-based diagnosis. Patient-specific pan-omics data analysis is going to disclose the genetic, epigenetic, and other functional profiles responsible for the disease of an individual which might eventually lead to development of individualistic ‘precision medicine’.

Cancer is a leading cause of death worldwide, being responsible for 9.6 million deaths till 2018 [2]. Cancer is a heterogeneous disease caused by aberrations of genes and proteins. ‘Precision oncology’ promises identification of disease subtypes, specific biomarkers and subsequently prediction and translation towards the development of treatment procedures. Pan-omics or multi-omics analysis in breast cancer has revealed significant differences in molecular subtype distribution [3]. Genomics and transcriptomics analysis of breast cancer data of Korean and Caucasian cohorts showed underlying molecular differences, which are responsible for the occurrence of breast cancer at the younger age in the Asian population compared to the western population [3]. Multi-omics analysis extended to different types of cancers confirms the existence of broadly two types of cancers, cancers caused by recurrent mutations and cancers caused by copy-number variations [4]. Machine learning based panomics analysis of pan-cancer data shows the existence of clusters within different types of cancers [5]. Machine learning based analysis has also been applied to identify cell-model selective anti-cancer drug targets for breast cancer [6].

Several data portals have been developed to make multi-omics data conveniently accessible. LinkedOmics contains pan-omics data of several types of cancers [7]. Databases like, GliomaDB [8] and MOBCdb [9] are dedicated to integrate multi-omics data for specific type of cancers. R packages like mixOmics [10] are available for the integration of multi-omics data. OmicsNet provides a web-based platform to create different types of molecular interaction networks for single or multiple types of omics data [11]. Network-based integration of multi-omics data allows to predict subnetworks of molecular interactions within a single type or multiple types of omics data [12]. R package Miodin [13] provides a software infrastructure for vertical and horizontal integration of multi-omics data but lacks a comprehensive network analysis and visualization. PaintOmics allows integrated visualization of multiple types of omics data in KEGG pathway diagrams [14]. Software package, MultiOmics Factor Analysis (MOFA) [15] integrates omics data in an unsupervised approach implementing generalized principal component analysis (PCA). pathfindR [16] finds active sub networks for genes in omics data and perform pathway enrichment analysis. R package Mergeomics [17] provides a pipeline to identify important pathways and key drivers in biological systems.

Different types of omics data carry information on different types of molecules, e.g. proteins, miRNAs, metabolites, etc. Hence, integrative analysis of pan-omics data needs a meta-interactome consisting of a protein-protein interaction network (PPIN) as well as different regulatory networks. The web server for the Analysis of Pan-Omics Data in Human Interactome Network (APODHIN) provides a unique platform where users can analyze different types of omics data using a human cellular meta-interactome network. This meta-interactome is the superimposition of regulatory networks of transcription factors (TFs) and miRNAs on PPIN. Graph theory based network analysis has become an essential tool for analysis of protein-protein interaction networks [18]. Centrality analysis measures the centrality of proteins in a network reflecting its importance in the construction and information flow of the network [19]. Over the years, different types of centrality measures have been used to find out insightful views of the network [20, 21]. In APODHIN, in addition to the creation of differentially omics data mapped meta-interaction network, we provide options to identify topologically important nodes (TINs) such as hubs, bottlenecks, and central nodes and their subsequent modules via protein-protein interaction (PPI) and regulatory relationship network analyses and pathway enrichment analysis. Important interacting nodes (IINs) (proteins and miRNAs) are identified based on overlap of TINs. Further, option is provided to correlate the potential of IINs/TINs to be prospective diagnostic and/or prognostic biomarkers. APODHIN also analyze and compare multiple omics data set for a single omics layer, such as transcriptomics, proteomics data collected from different patient cohorts and/or different stage/grade of the same cohort.

Additionally, APODHIN utilizes multi-omics data and calculate cross-pathway regulatory and PPI links connecting signaling proteins or transcription factors or miRNAs to metabolic enzymes and their metabolites using network analysis and mathematical modeling. These cross-pathway links were shown to play important roles in metabolic reprogramming in cancer scenarios such as glioblastoma multiforme in a previous work [22].

In addition to the server part, APODHIN also has a database part where TINs and cross-pathway links were identified using publicly available omics datasets collected for various gynecological cancers. Dataset specific and common TINs and cross-pathway links were provided in the APODHIN database part. APODHIN platform is freely available at http://www.hpppi.iicb.res.in/APODHIN/home.html.

## Materials and methods

### Server description

APODHIN web server is dedicated for the integration and subsequent analysis using single or multiple types of omics data. For single type of omics data, APODHIN can analyze multiple datasets (up to 3) which may correspond to either different stages of a disease from a single cohort or from dataset collected from multiple patient cohorts and/or cell lines.

For multiple types of omics data, APODHIN allows single input data file for each type of omics data. Following sections briefly describe the various analytical part of the APODHIN server.

### Data collection

APODHIN web server is preloaded with a human cellular meta-interactome network. This meta-interactome consists of human protein-protein interaction network (HPPIN), network of human miRNAs and their target genes and network of human transcription factors (TFs) and their target genes. The protein-protein interaction data was collected from STRING [23] database (version 11). Interactions having an experimental score ≥ 700 are only considered. This extracted and processed HPPIN consists of 34405 interactions from 15164 proteins. The degree distribution of this network is found to be scale-free and follows the Power law, *P(k)* ~ *k*^*-γ*^ where *γ* is 1.9 [18, 24]. Target gene information of miRNAs was collected from the TarBase [25] and miRTarBase [26] databases. From the TarBase database (version 6) we have taken interactions supported only by reliable low-throughput experimental data whereas miRNA target interactions with strong confidence from miRTarBase (version 6) were considered for APODHIN meta-interactome network. We found 2492 target genes for 544 miRNAs creating 6917 interactions. TFs and their target genes were downloaded from Human Transcriptional Regulation Interactions database (HTRIdb) [27]. We found 11887 target genes for 284 TFs creating 18153 interactions. These three networks were merged together to form the APODHIN meta-interactome consisting of two types of biomolecular nodes i.e., proteins/genes and miRNAs along with three types of interactions, i.e, protein-protein, miRNA-target gene, and TF-target gene, respectively.

Additionally, we have also included a network of metabolites as substrate and product with their corresponding metabolic enzymes in the APODHIN server. For constructing this network, we downloaded metabolic reactions from MetaNetX database [28] and extracted the metabolites along with the corresponding metabolic enzymes and further filtered those enzymes and metabolites which have been listed in the Human Metabolome Database (HMDB) database [29].

### Pan-omics data integration and meta-interaction network extraction

In APODHIN web server, user can upload single or multiple types of omics data. The server accepts RNA transcriptomics, miRNA transcriptomics, proteomics, phosphoproteomics, genomics, epigenomics and metabolomics data. The current version of the server accept only processed format of the omics data where differential expression/abundance of corresponding biomolecules are provided with *log*FC for defining up and down regulation of genes/miRNAs/proteins and threshold probability or *p*-value. For RNA transcriptomics, miRNA transcriptomics and proteomics data user should select threshold values of *log*FC for defining up and down regulation of genes/miRNAs/proteins and corresponding adjusted *p*-value. Uploaded files should contain list of genes/miRNAs/proteins along with *log*FC and *p* values. Sample file formats for different omics data are provided in the APODHIN help page. For genomics, epigenomics, and phosphoproteomics data, genes that are mutated and/or methylated and proteins, which are phosphorylated are considered, respectively.

APODHIN web server extracts the interactome networks from the parent meta-interactome for the genes, mRNAs, miRNAs, proteins, and metabolites that are either deregulated or altered according to the user supplied single or multiple omics data. It creates a filtered meta-interactome network comprising of deregulated or altered nodes and their 1st or 2nd level (as chosen by user) interactors and/or regulators. For metabolomics data, the web server finds out the proteins linked with metabolites and constructs network. These single or multi omics data specific meta-interactome networks are subsequently displayed in an interactive three-dimensional (3D) network viewer within the APODHIN server.

For the module ‘pathway connectivity analysis’, RNA transcriptomics, miRNA transcriptomics, and proteomics data were considered as primary data and submission of at least one of them is mandatory to define deregulated miRNAs and/or genes/proteins. In case of ‘pathway connectivity analysis’, the *log*FC values for each of the uploaded omics data is normalized in the scale of −1 to +1 where positive and negative values indicate up and down regulated entities, respectively. If more than one primary omics data, for example, transcriptomics and proteomics are provided, APODHIN web server sums up the normalized logFC values from the different omics data for the same node (RNA/protein) and if the sum is non-zero, gene/protein is considered deregulated. Details of the utilization of the normalized omics values in mathematical modeling based pathway connectivity link identification are provided later. In this module, the information on metabolites for any enzyme can be obtained in the associated table on selection of enzyme.

### Network analysis and identification of TINs

Once the context specific meta-interactome network is formed via utilization of user supplied single or multiple omics data, APODHIN web server primarily finds three types of topologically important nodes (TIN), namely, hubs, central nodes (CNs) [30] and bottlenecks (BNs) [31]. To find the important nodes, network and node indices like degree, betweenness, closeness and clustering coefficients are calculated from the extracted meta-interactome network. These nodes were calculated using previously reported methods and protocols [30]. For transcriptomics and proteomics data, TINs are identified from the expressed nodes only. For phosphoproteomics, genomics, epigenomics and metabolomics data, TINs are identified from phosphorylated, mutated, methylated proteins/genes and metabolic enzymes, respectively.

Hubs are nodes that have high degrees. APODHIN web server converts normalized degree distribution to corresponding z-score distribution. The plot of probability distribution function (PDF) of z-scores for all nodes in network is sent to the user by email. Based on the user provided threshold value of degree, hub nodes are identified. Scores concerning individual centrality parameters like, betweenness, closeness and clustering coefficients are calculated and the cumulative centrality scores (CCS) are estimated by summing over the combined scores for first layer interactors. Normalized cumulative centrality scores are converted into z-scores. The PDF of z-scores for all nodes of network are sent to the user by email and CNs are chosen based on the user provided threshold value of z-score [30]. Bottleneck nodes are characterized based on their betweenness values. Normalized betweenness values are converted into z-scores. Similar to CNs, bottleneck nodes are also chosen based on the user provided threshold z-score, chosen from the PDF plot of z-score for all nodes.

Further, sub-network consisting of TINs and their first and second layer interactors are constructed and displayed in an interactive three-dimensional (3D) network viewer.

### Pathway mapping and network of mapped pathways

For each identified TIN, particularly for genes and proteins, APODHIN maps the corresponding pathways listed in the KEGG database [32]. APODHIN performs a hypergeometric Fishers Exact test and selects enriched pathways satisfying *p*-value (*p*_*HGD*_) ≤ using the following contingency table and formula.

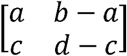

Where, a = Number of genes in the pathway

b = Number of genes in the gene list

c = Total number of genes in the pathway

d = Total number of genes in all pathways in KEGG

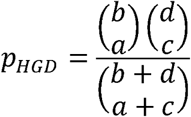

Further, a network representation of important nodes along with their enriched mapped pathways is displayed in an interactive three-dimensional (3D) network viewer. Figure 1A shows the flow chart of ‘pan-omics data mapping and network analysis’ module of APODHIN.

**Figure 1:**
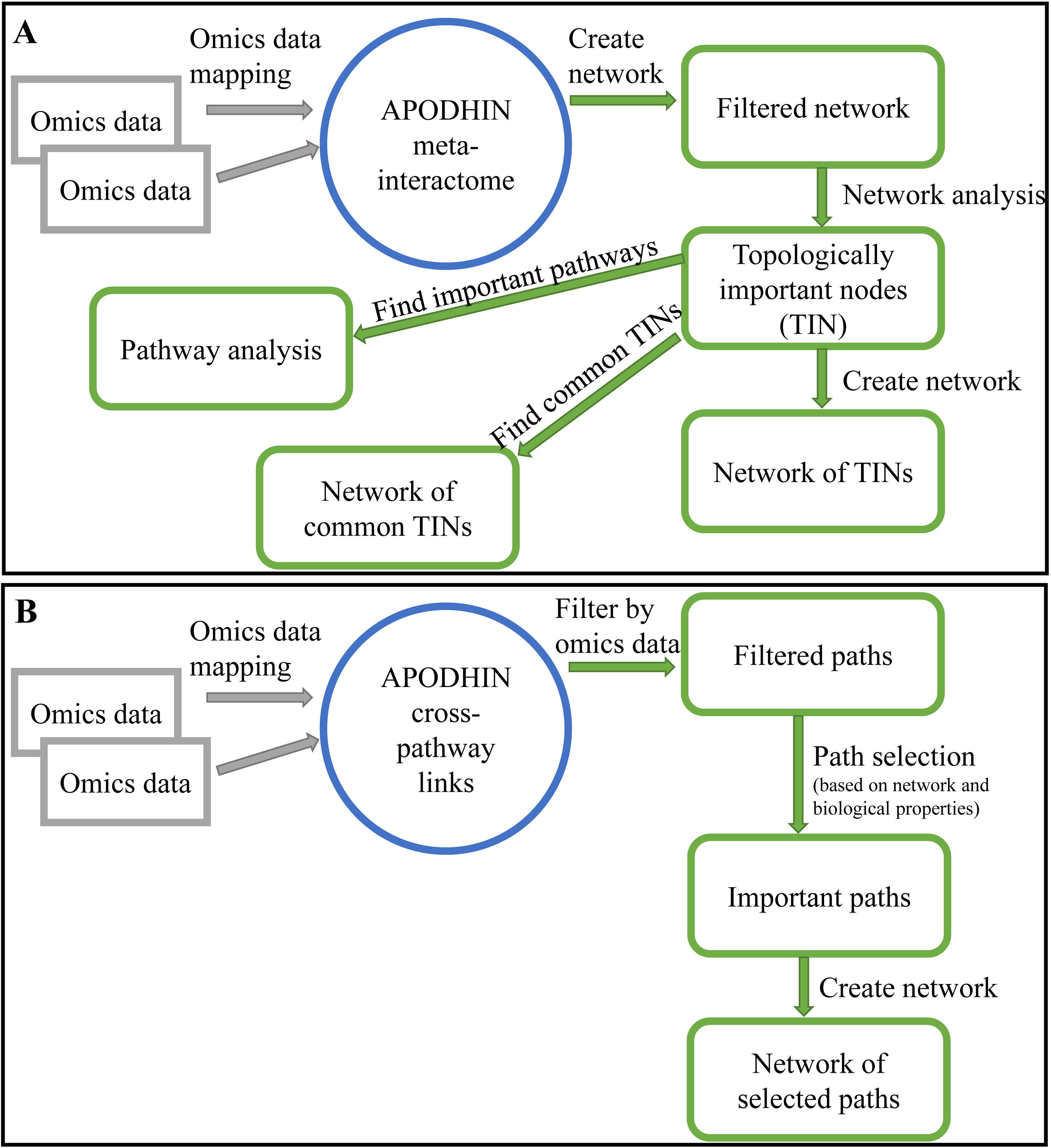
Flow charts showing work flow in APODHIN web-server for module (A) data mapping and network analysis and (B) pathway connectivity analysis.

### Pathway connectivity analysis and cross-pathway links

This module of the APODHIN web server aims to construct regulatory interaction networks and subsequently identifies cross-pathway interaction links connecting different cellular pathway proteins (e.g., signaling proteins (S)), regulatory proteins (e.g., transcription factor (TF)) or miRNAs with metabolic pathway proteins (M).

For this purpose, APODHIN web server was preloaded with cross-pathway links or paths where protein-protein interactors (P) connect X nodes (X can be S or target gene of TF or target genes of miRNAs) with M (metabolic) proteins. We have limited the number (n) of protein-protein interactors (P) to a maximum value of three between X and M proteins. This limit provides four types of paths, XM (*n* =0), XPM (*n* =1), XPPM (*n* =2), XPPPM (*n* =3). These cross-pathway linking paths are filtered and selected based on expression and/or abundance status of the biomolecules supplied by user uploaded pan-omics data for a given disease or context. The filtering criteria for any given path is set when the terminal nodes are found to be deregulated and the remaining nodes are at least expressed within the user provided single or multi-omics datasets.

We implemented an established probabilistic approach based on the Hidden Markov Model (HMM) [22, 33, 34] utilizing the information of experimentally established protein-protein interactions and gene regulatory information to extract novel paths and interconnections between regulatory nodes such as signaling proteins, transcription factors and miRNAs and metabolic pathway proteins (M). Within these important X-M pairs, important cross-pathway connecting paths are again scored by considering all filtered paths between X-M pairs. To find important X-M pairs, weights are assigned on nodes and edges depending on network and biological properties. Edge weight is assigned in terms of normalized interaction probability which is proportional to the product of their expression scores.

Two types of node weights, network entropy, and effect-on-nodes are considered. Network entropy includes local entropy of the node. Another node weight parameter, ‘effect-on-node’ considers the impact of interactors of a particular gene in the cross-connected network. The ‘effect-on-node’ considers both biological and network properties of the node. Biological properties include deregulated gene, signaling crosstalk gene and rate limiting enzyme. Network properties include hubs, CNs, and bottlenecks.

APODHIN web server allows the user to choose maximum four weight options out of the six weights. If a node satisfies any of the selected weight options, weight value 1 is assigned for each satisfied option. To identify important cross-connecting X-M pairs we have evaluated ‘path score’ (PS) based on a Hidden Markov Model (HMM) implemented within the core mathematical model that calculated the significant cross-pathway linking paths. ‘Path scores’ are converted to z-scores and paths having z-score 1 are considered as important cross-connecting paths. A detailed description of the mathematical models and path calculation is available in our previous publication [22]. Figure 1B shows the flow chart of ‘pathway connectivity analysis’ module of APODHIN.

### APODHIN architecture

APODHIN web server is created using HTML, PHP, PYTHON, and JAVA scripts. Client/user side scripts are written in HTML, PHP and JAVA scripts. User uploaded data is analyzed using PYTHON scripts. For network analysis, PYTHON package networkX (version 1.8.1) is used. For visualization of 3D presentation of networks JAVA scripts based open source technologies (*three.js* and *3d-force-graph.js*) were utilized.

APODHIN has two separate parts A. APODHIN server and B. APODHIN database.

### APODHIN server

APODHIN web server is preloaded with human interactome network containing PPIN, target gene network of miRNAs and target gene network TFs. Proteins participating in signaling and metabolic pathways are also marked separately. Metabolites along with their target enzymes are also included within APODHIN. This meta-interactome network is used as framework of cellular interactions and is further used to map user supplied single or multiple types of ‘omics’ data to perform the following analyses.

- Omics data mapping and network analysis: This module has two sub-modules. On clicking first submit button, this web server provides meta-interactome network filtered by uploaded omics data where deregulated and/or altered nodes along with their interactors are included. Users can further proceed for finding important interacting nodes from the ‘pan-omics’ data mapped interaction network by clicking second submit button. Tabular, graphical and 3D network representations of the identified TINs are provided for better appreciation. Overlap of the TINs is shown both in tabular and interactive 3D network visualization. Additionally, TINs and their enriched pathways are also shown in tabular and interactive 3D network visualization manner. Sample input files for each omics data type and example analysis output are provided for the ease of use and apprehension.
- Pathway connectivity analysis: As mentioned before, this sub-module highlights significant protein-protein interaction and regulatory paths connecting signaling proteins/transcription factors (TF)/miRNAs to metabolic proteins. These cross-pathway links are thought to be supra-molecular regulatory links/signatures connected with metabolic rearrangement or reprogramming events that are observed during cancer. In APODHIN, these cross-pathway regulatory links can be constructed from three types of interaction networks.

1. Integrated network where signaling (S) and metabolic (M) pathway proteins are connected through protein-protein interactors (P).
2. Integrated network where target genes of TFs and metabolic (M) pathway proteins are connected through protein-protein interactors (P).
3. Integrated network where miRNA target genes and metabolic (M) pathway proteins are connected through protein-protein interactors (P).

Cross-pathway linking paths are filtered and selected based on expression and/or abundance status of the biomolecules supplied by user uploaded single or pan-omics data for a given disease or context. These paths are shown both in tabular and interactive 3D network visualization.

### APODHIN database

We have used the APODHIN web server to construct individual cancer and dataset centric meta-interactome network using cell line specific single and/or multi-omics data collected from various resources such as GEO [35], PRIDE [36], publication reports and data sources for cervical, ovarian, and breast cancers, respectively. Further, these cancer and dataset specific meta-interactome networks were analyzed and important interacting nodes and cross-pathway links were identified and provided within the APODHIN database module. We have used cancer cell line derived omics data freely available from different public resources. Options are provided for the users to select single and/or multi-omics data to construct the meta-interactome networks and further analyze them to identify and important interacting nodes and cross-pathway links specific for the selected dataset.

## Results

### Input options

APODHIN server provides two different but linked analysis options for the users who would like to utilize single or multiple types of omics data for a given context. Users can construct meta-interaction networks related to the altered and/or deregulated biomolecules, say DNA, RNA, proteins, and metabolites, respectively via utilization of specific data types singly or in combination. Further, these context (disease) specific meta-interactome networks are utilized to identify important interacting nodes comprising regulatory entities such as miRNA or transcription factor and effector entities like interacting proteins or enzymes. Specific TINs and their related regulatory, PPI, or enzyme-metabolite interactions can be crucial for mechanistic understanding of the complex systemic diseases like cancers. Hence, APODHIN web server provides options to upload seven types of ‘omics’ data comprising of mRNA transcriptomics, miRNA transcriptomics, proteomics, phosphoproteomics, genomics, epigenomics, and metabolomics. The file formats for each data type is specified in the ‘Help’ page and sample input files are also available in the server input page. For transcriptomics and proteomics data, maximum and minimum threshold values for the differential expression/abundance (*log*FC) and statistical significance of that (*p*-values) need to be provided. As the calculations are computation intensive, results are sent via email.

Similarly, for cross-pathway connectivity analysis users need to upload single or multiple types of ‘omics’ data for a given context. Now, in this case, users also need to specify the type of connectivity they would like to explore, for example, signaling to metabolic proteins, TFs to metabolic proteins, or miRNAs to metabolic proteins. Only one type of pathway connectivity can be explored at a time for a given set of ‘omics’ data. Additionally, users also need to select the kind of weights (see Methods) that would be applied while calculating the scores of the selected cross-pathway regulatory and PPI paths. E-mail address needs to be supplied for APODHIN server to send the result link of the identified cross-pathway connections.

### Output options

Output option for the ‘Data mapping and network analysis’ module has two stages. At first stage (Figure 2A), the context specific meta-interactome network (‘filtered network’) can be visualized via a user interactive 3D network viewer where information regarding each node and edge are provided in graphical as well as tabular view (Figure 2B). Status of the ‘omics’ data mapping is shown in various color codes for the nodes whereas different relationship like protein-protein interaction, miRNA-target gene interaction, and TF and target in connections are shown varied color codes. Additional details about the protein nodes can be obtained via GeneCards [37] link while miRNA details can be found via miRTarBase [26] link. List of metabolites mapped onto the protein nodes are also provided both in the network viewer as well as in the adjacent tabular format. If network analysis is opted, along with filtered network, APODHIN provides the probability distribution functions for the opted TINs (Figure 2A). After receiving the user provided threshold values, APODHIN finds TINs and provides a tabular result with a summary of the user uploaded data (Figure 2C). Filtered nodes (genes/proteins/miRNAs) that are satisfied the selected threshold criteria are further utilized for meta-interactome network construction. Users can see the number of TINs (as hub, bottlenecks and central nodes) and their mutual overlap using interactive Venn diagrams by clicking the ‘link for analysis’ option for single or combination of ‘omics’ data, The resultant page (Figure 2D) provides three output options. Combined analysis of multiple types of files is shown if multiple types of files are selected. Here also, the resultant page (Figure 2E) provides three output options. First, the regulatory and PPI connectivity specific to the hubs, bottleneck and central nodes can be seen via corresponding link where networks of deregulated hubs, bottleneck, and central nodes can be seen separately and saved accordingly (Figure 2F). Association to various kinds of cancers for the identified TINs as favorable/unfavorable prognostic markers are also provided here after mapping the TINs to the data provided in Human Proteome Atlas [38]. Another option provides the network of common TINs (Figure 2G) whereas a separate link provides network of enriched pathways with the identified TINs (Figure 2H). Enriched pathway networks of deregulated hubs, bottleneck, and central nodes can be seen separately and saved accordingly. In all these three network output options, data can be downloaded in text format for further analysis.

**Figure 2:**
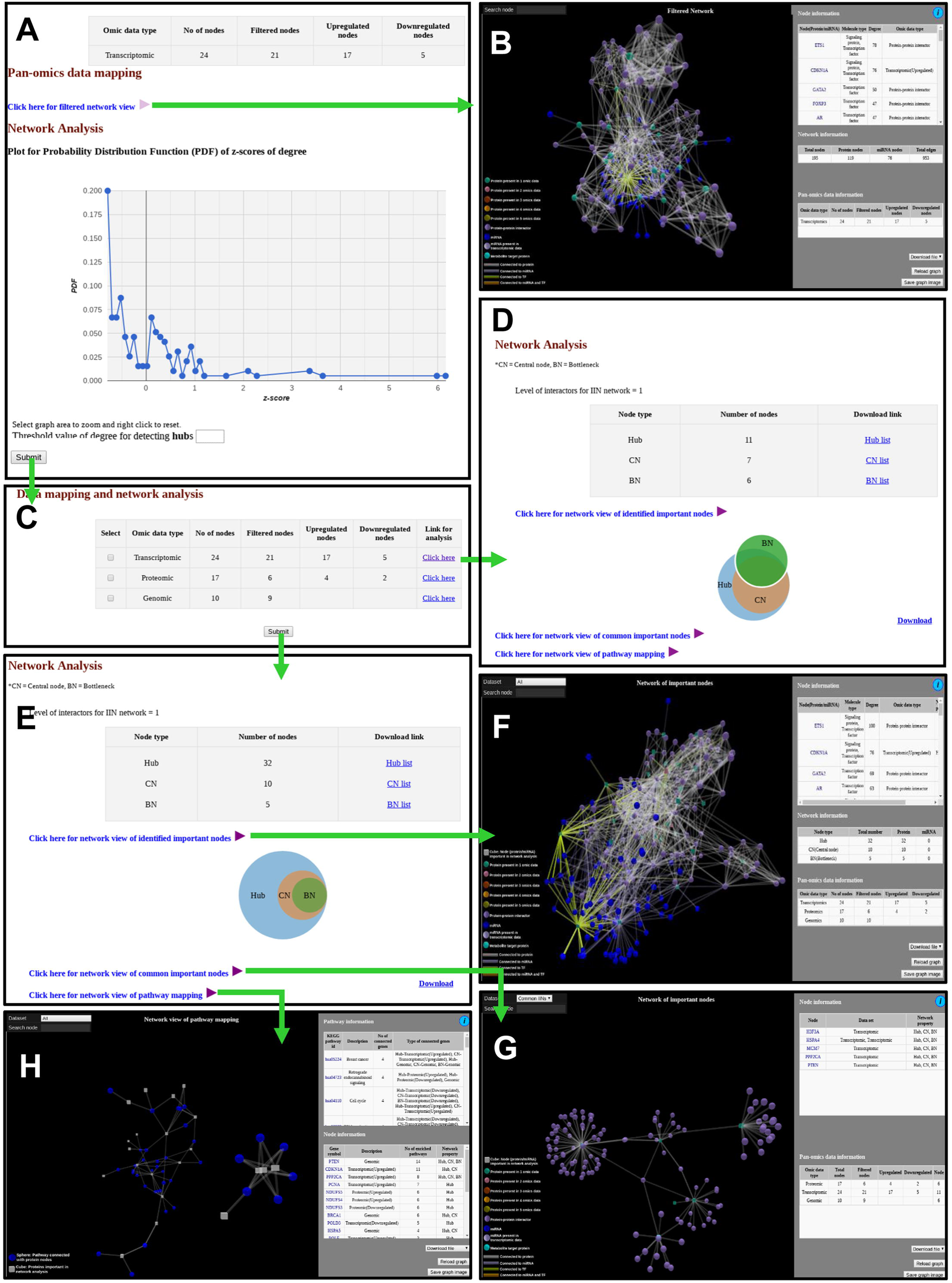
Snapshots of outputs of module ‘data mapping and network analysis.’ (A) Page showing link for filtered network and probability distribution function. (B) Filtered network. (C) User provided input data in tabular form along with link for analysis for single omics data, shown when analysis is over. (D) Output page of a single omics data. (E) Network analysis page for multi-omics data. (F) Network of important interacting nodes. (G) Network of important nodes. (H) Network of pathway mapping.

Similar to ‘Data mapping and network analysis’, ‘Pathway connectivity analysis’ module also provides a tabular result with a summary of the user uploaded data (Figure 3A). Users can see the cross-pathway links for single (Figure 3B) or combination (Figure 3C, 3D) of ‘omics’ data in the 3D network visualization window where significant protein-protein interaction and regulatory paths connecting signaling proteins/TFs/miRNAs to metabolic proteins are shown in color coded fashion. As described before, these cross-pathway links or paths connect X nodes (X can be S or target gene of TF or target genes of miRNAs) with metabolic (M) proteins. These linking paths are filtered and selected based on expression and/or abundance status of the biomolecules supplied by the users where for any given path the terminal nodes are found to be deregulated and the remaining nodes are at least expressed. The corresponding pathways and biological functions of the proteins are also provided in tabular format adjacent to the network viewer. Additionally, the metabolites connected to the metabolic proteins that are part of the selected cross-pathway links are also provided in the same page.

**Figure 3:**
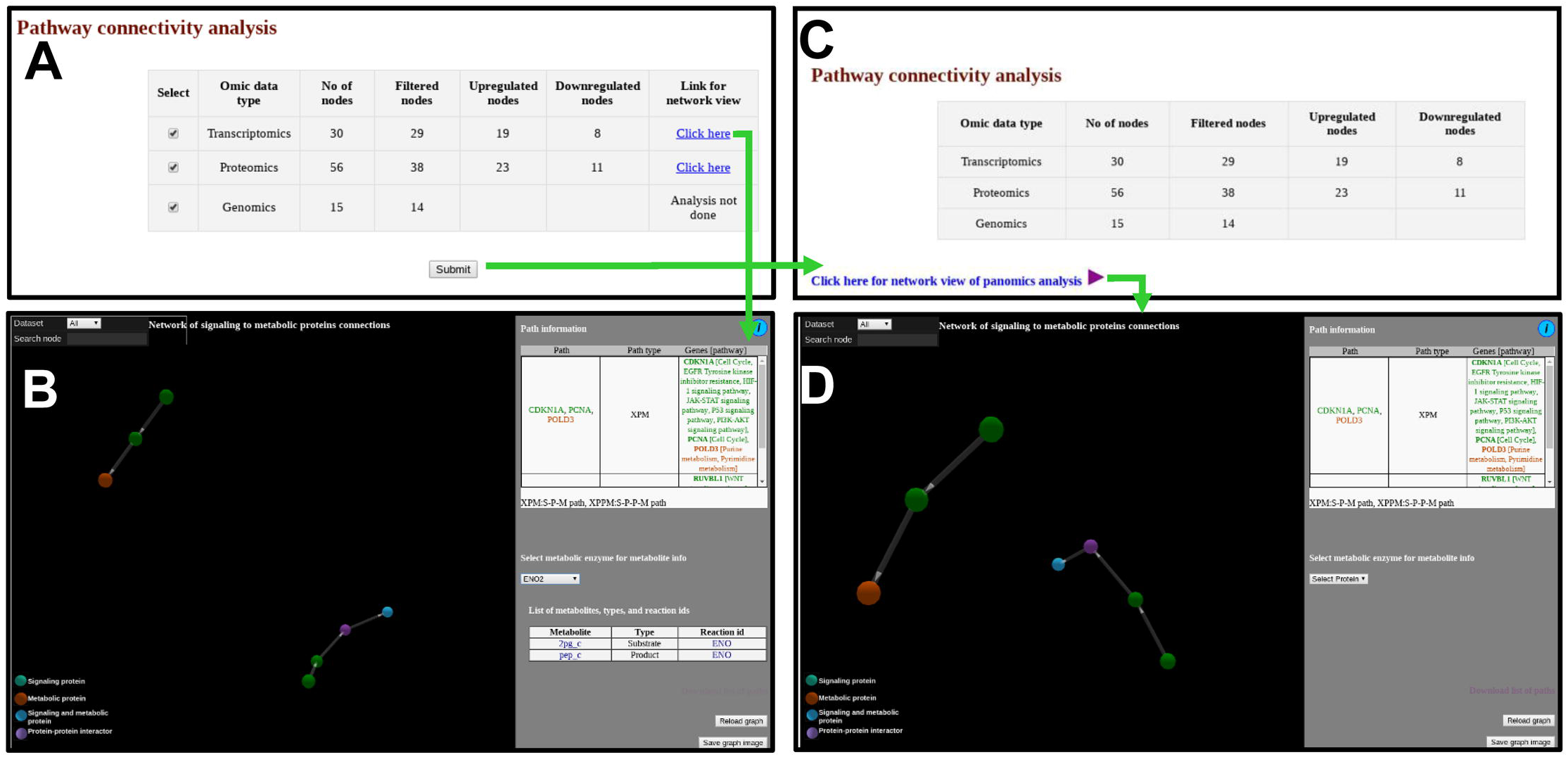
Snapshots of outputs of module ‘pathway connectivity analysis.’ (A) Output page shows user provided data in tabular form along with link for analysis for single omic data. (B) Output page showing network of signaling to metabolic proteins connecting paths for a single type of omics data. (C) Output page when multiple types of omics data is selected in (A). (D) Network of signaling to metabolic proteins connecting paths for pan-omics data.

### Database option

APODHIN database contains important nodes (genes/proteins/miRNAs), pathways, and their networks with interacting partners specific for cancers affecting women such as cervical, ovarian, and breast cancer. This database also contains important paths linking signaling proteins/TFs/miRNAs to metabolic enzymes, which could perhaps be responsible for metabolic reprogramming in cancer. The database content is produced by APODHIN web server using publicly available cervical, ovarian, and breast cancer specific cell line based omics data. Figure 4 briefs the statistics derived from APODHIN database for mRNA transcriptomics data of different cell lines of cervical, ovarian and breast cancer. Panels A and B show the overlap of up-regulated and down-regulated genes, respectively. It reveals lesser overlap among deregulated genes across cell lines for all cancers. Nodes satisfying any two types of TINs are considered as important interacting nodes (IINs). Panel C shows overlap for common IINs where higher overlaps between cell lines across cancer types are observed. Similarly, panel D shows much higher overlap of common pathways mapped by IINs. This demonstrates that IINs and their pathways represent the common core genes and processes related to a cancer type in a better way than that achieved by the initial deregulated genes obtained from the omics data. Panel E and F show the number and overlap of deregulated genes and IINs as prognostic markers of respective cancer type. Figure S1 denotes higher fraction of prognostic markers among the IINs and TINs (except BNs) compared to the deregulated genes.

**Figure 4:**
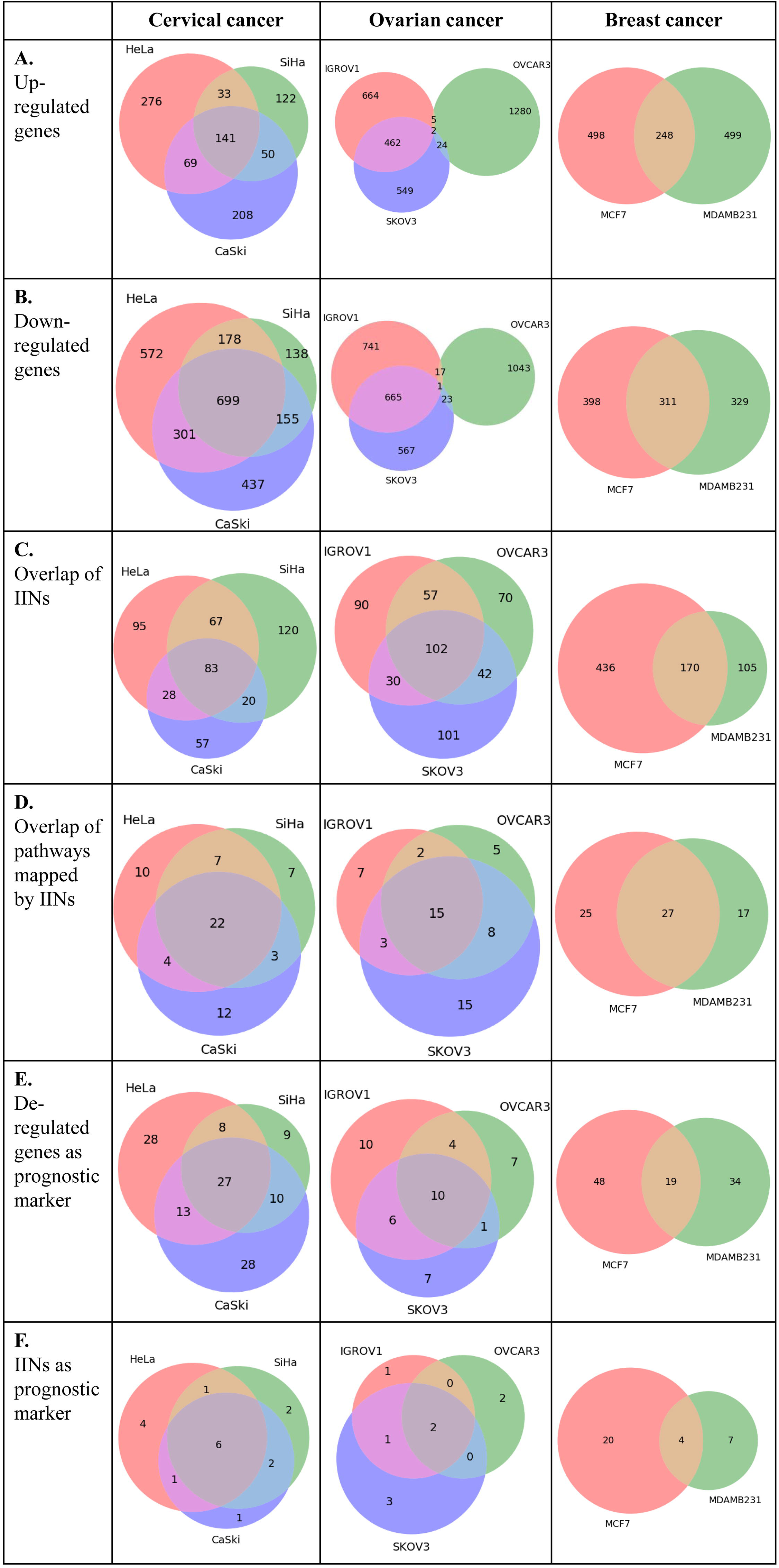
Statistics of APODHIN database features using mRNA transcriptomics data derived from cervical (HeLa, SiHa, and CaSki), ovarian (IGROV1, SKOV3, OVCAR3) and breast cancer (MCF7 and MDAMB231) cell lines. Transcriptomics data was derived from the GEO datasets GSE9750, GSE19352, and GSE71363, respectively.

Figure 5 shows overlap of cross-pathway links or paths connecting signaling (S) proteins, TFs, and miRNAs to metabolic (M) proteins identified using omics data derived from the cell lines of three types of cancers. For signaling to metabolic connection, 13 common paths including 2 common paths for three cervical cell lines were observed. However, no such overlap was found for breast and ovarian cancer cell lines.

**Figure 5:**
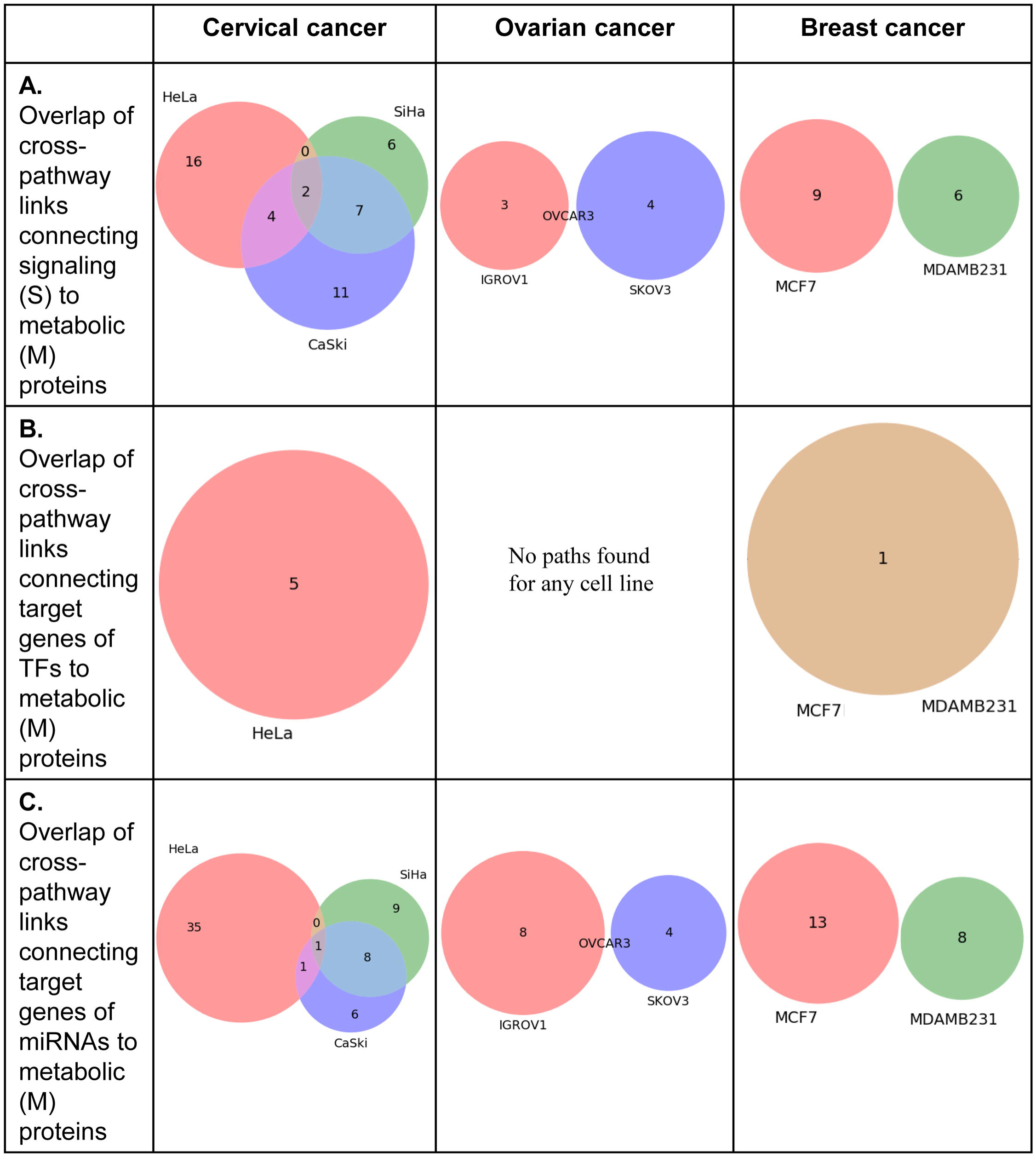
Statistics of ‘pathway connectivity analysis’ module of APODHIN database using mRNA transcriptomics data derived from cervical (HeLa, SiHa, and CaSki), ovarian (IGROV1, SKOV3, OVCAR3) and breast cancer (MCF7 and MDAMB231) cell lines. Transcriptomics data was derived from the GEO datasets GSE9750, GSE19352, and GSE71363, respectively.

## Discussion

Large-scale genomics, transcriptomics and proteomics approaches have made it possible to characterize different clinical spectra associated with cancers. Use of pan-omics platforms and approaches in the analysis of systemic disease like cancer will not only help to identify numerous useful biomarkers but also will expose areas for further improvement in therapeutic intervention. Here, we present APODHIN web server, which extracts cellular interactome networks from the parent meta-interactome for the genes, mRNAs, miRNA, proteins, and metabolites that are either deregulated or altered according to the user supplied single or multiple omics data. These single or multi-omics data specific meta-interactome networks are utilized to identify TINs and their sub-modules enriched with protein-protein interaction and regulatory relationship via utilization of graph theory based network analyses and biological pathway enrichment analysis. Important interacting nodes (proteins and miRNAs) are identified based on overlap of keys nodes such as hubs and bottlenecks which could correlate to be prospective diagnostic and/or prognostic biomarkers or even turn out to be potential therapeutic targets.

Molecular mechanisms for cancer progression and development of potential therapeutics to inhibit these complex diseases are difficult from the independent knowledge of signaling, transcription factors, miRNAs, and metabolic pathways. Metabolic reprogramming is an essential hallmark of cancer [39]. Understanding the coordination among various cellular pathways, such as gene-regulatory, signaling and metabolic pathways is crucial and may provide clues into the molecular mechanism of metabolic adaptation in cancer and associated cells. Therefore, there is an urgent need for systems biology model, which can coordinate among signaling-induced proliferation of tumor cells/growth, transcription factor/miRNA based gene regulation and metabolic processes. Hence, we emphasized to design a mathematical approach to identify significant proteins forming interconnections between signaling, regulatory and metabolic pathways. We have constructed an integrated network where signaling (S), regulatory (TFs and miRNAs) and metabolic (M) pathway entities are connected through protein-protein and gene regulatory interactions. Interconnections between regulatory components such as signaling proteins/TFs/miRNAs and metabolic pathways need to be elucidated rigorously to understand the role of oncogene and tumor suppressors in regulation of metabolism alongside their normal functions. Analyses of such cross-connected network and linking paths will facilitate probable way(s) to inhibit cancer progression in a more specific manner.

Considering the growing demand of multi-omics data integration followed by systems biology based analytical interpretation of the large-scale ‘omics’ data, implementation of a robust and user-friendly web-based platform is very much due. In order to make better sense out of the various ‘omics’ data, it is imperative to utilize them in a way so that the global scenario of the complex and multi-layer cellular interactome can be recapitulated. Several data portals have been coming up to make multi-omics data accessible, visible and more importantly, interpretable. Web-servers like OmicsNet [11] is a technically powerful web based platform specifically meant for better visualization of molecular networks. It mainly provides varied and efficient ways of network visualization including different components. However, it provides minimal emphasis on networks analysis and identification and interpretation of important interacting nodes and cross-pathway links. Similarly, this server only accept differential omics data for genes/proteins and metabolites, it does not have the option to include the epigenetic modification, miRNA expression data, and phosphoproteomics data. APODHIN is perhaps the only available web based platform that offers to a) integrate multi-omics data onto an exhaustive multi layered cellular meta-interactome network, b) extract and analyze the context specific networks and sub-networks to identify TINs that could serve as potential biomarkers and/or therapeutic targets c) rationalize the role of the identified TINs to the given context via pathway enrichment and prognostic marker correlation, and d) identify cross-pathway interconnections between regulatory components such as signaling proteins/TFs/miRNAs and metabolic pathways for better understanding the role of oncogenes and tumor suppressors in regulation of metabolic reprogramming during cancer. IINs and TINs identified by the APODHIN shows higher fraction of important prognostic marker indicating the usefulness of the TINs/IINs over only deregulated genes. Table 1 provides a qualitative comparison of features and functionalities of APODHIN with respect to existing omics data analysis tools.

**Table 1:** Comparison of APODHIN features and functionalities with existing pan-omics data analysis tools.

| Feature | APODHIN | OmicsNet [11] | mixOmics [10] | iOmicsPASS [12] | Miodin [13] |
| --- | --- | --- | --- | --- | --- |
| Platform | Web | Web | Standalone | Standalone | Standalone |
| Programming language | Python, R, perl | R | R | C++ | R |
| Types of omics data as input |  |  |  |  |  |
| mRNA transcriptomics | Yes | Yes | Yes | Yes | Yes |
| miRNA transcriptomics | Yes | Yes | Yes | No | Yes |
| Proteomics | Yes | Yes | Yes | Yes | Yes |
| Phospho-proteomics | Yes | No | Yes | No | No |
| Genomics | Yes | Yes | Yes | Yes | Yes |
| Epi-genomics | Yes | No | No | No | Yes |
| Metabolomics | Yes | Yes | Yes | No | No |
| Multiple lists of same type of omics data | Yes | No | Yes | No | Yes |
| Finding deregulated proteins/genes/miRNAs | Yes | No | No | No | No |
| Map in meta-interactome | Yes | No | No | No | No |
| 3D interactive network | Yes | Yes | No | No | No |
| Network topology analysis | Yes | No | No | No | No |
| Prognostic status of proteins/genes (in cancer) | Yes | No | No | No | No |
| Pathway enrichment analysis | Yes | Yes | No | Yes | No |
| Analysis for regulatory network protein links | Yes | No | No | No | No |

However, there is still scope for improvement for the APODHIN server and database. For example, in future we would like to equip the server to accept and process raw ‘omics’ data directly and further create the processed data for genetic or epigenetic alterations, differential expression and abundance, respectively. Similarly, the server and the database should be enriched in such a way that it could be utilized for deep learning and artificial intelligence based tools to predict the disease outcome, recurrence and drug resistance, respectively.

## Authors’ contributions

SC conceptualized the project. NB and SC designed the web server. NB created the web server. KK, SB and RB provided data for meta-interactome network. NB and SC analyzed the data and drafted the manuscript.

## Acknowledgement

The authors acknowledge CSIR-Indian Institute of Chemical Biology for infrastructural support. SC acknowledges the Systems Medicine Cluster (SyMeC) grant (GAP357), Department of Biotechnology (DBT) for funding. NB acknowledges the Systems Medicine Cluster (SyMeC) grant (GAP357), Department of Biotechnology (DBT) for fellowship. KK, RB, and SB acknowledge Department of Biotechnology (DBT), Council of Scientific and Industrial Research (CSIR), respectively for their fellowships.

## Supplementary Figure

**Figure S1:** Comparison of fraction of prognostic markers within the deregulated genes and network analysis derived important nodes, such as IIN and various TIN (e.g., Hubs, CN, and BN, respectively).

