## Supplementary figures and images for "Analysis of Pan-Omics Data in Human Interactome Network (APODHIN)"

### Figure S1

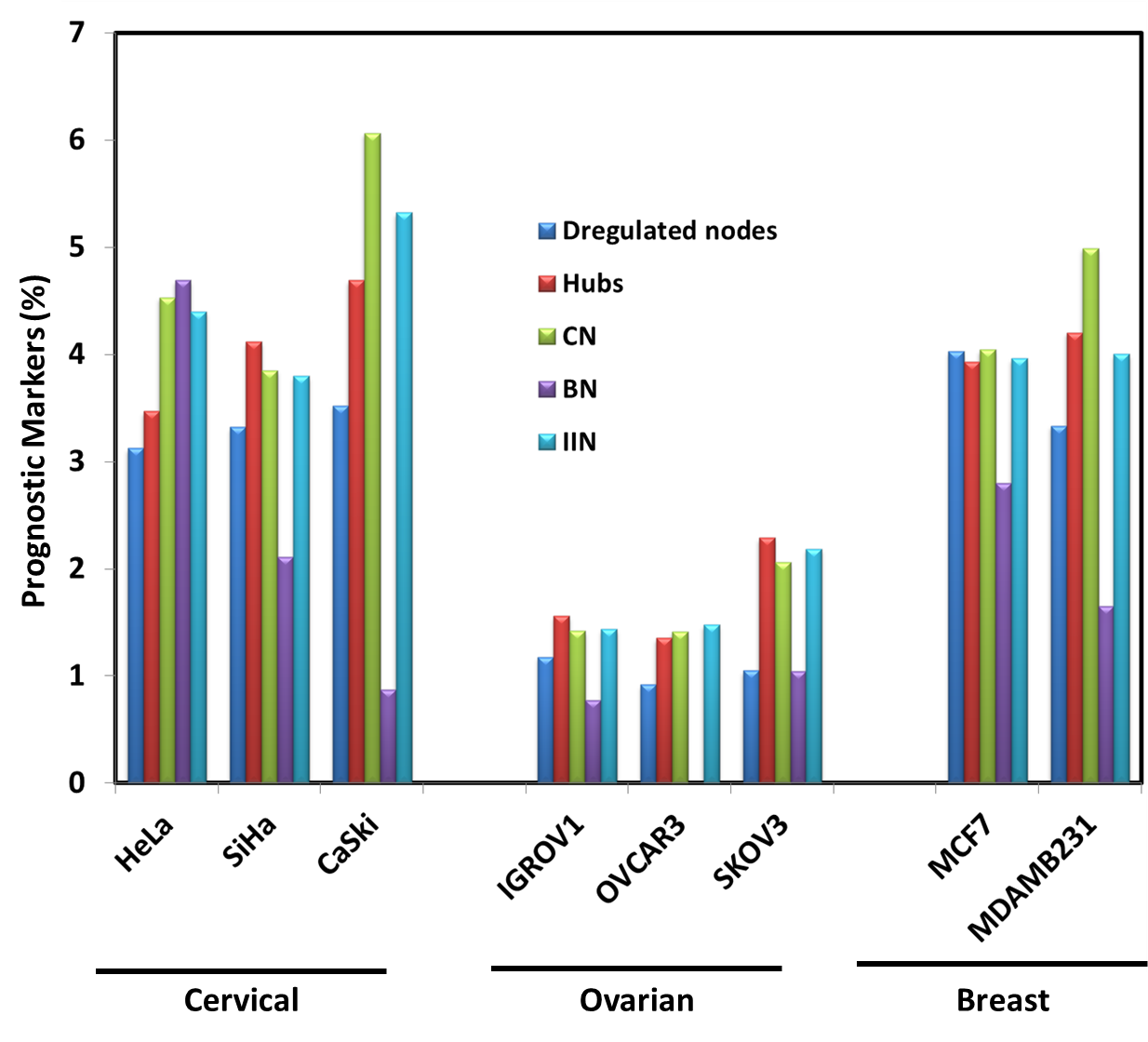
